# Precision inhibitory stimulation of individual-specific cortical hubs disrupts information processing in humans

**DOI:** 10.1101/254417

**Authors:** Charles J. Lynch, Andrew L. Breeden, Evan M. Gordon, Joseph B. C. Cherry, Peter E. Turkeltaub, Chandan J. Vaidya

**Affiliations:** Department of Psychology, Georgetown University; VISN 17 Center of Excellence for Research on Returning War Veterans; Department of Psychology and Neuroscience, Baylor University; Center for Vital Longevity, School of Behavioral and Brain Sciences, University of Texas at Dallas; Neurology Department, Georgetown University Medical Center; Research Division, MedStar National Rehabilitation Hospital; Children’s National Medical Center, Washington DC

**Author notes:** Correspondence (C.J.L.), (C.J.V.). Author contributions: CJL, ABL, EMG, PET and CJV designed the experiment. EMG contributed key analytic tools. CJL and JBBC performed research. CJL, ALB, EMG processed and analyzed the data. CJL, ALB, EMG, PET, CJV wrote the manuscript with input from all authors.

**Keywords:** Brain stimulation, connectomics, hubs, resting-state fMRI

## Abstract

Non-invasive brain stimulation (NIBS) is a promising treatment for psychiatric and neurologic conditions, but outcomes are variable across treated individuals. This variability may be due in part to uncertainty in the selection of the stimulation site – a challenge complicated further by the variable organization of individual human brains. In principle, precise targeting of individual-specific brain areas serving outsized roles in cognition could improve the efficacy of NIBS. Network theory predicts that the importance of a node in network can be inferred from its connections; as such, we hypothesized that targeting individual-specific “hub” brain areas with NIBS would impact cognition more than non-hub brain areas. We first demonstrate that the spatial positioning of hubs is variable across individuals, but highly-reproducible when mapped with sufficient per-individual rsfMRI data. We then tested our hypothesis in healthy individuals using a prospective, within-subject, double-blind design. We found that inhibiting a hub with NIBS disrupted information processing during working-memory to a greater extent than inhibiting a non-hub area of the same gyrus. Furthermore, inhibition of hubs linking specific control networks and sensorimotor systems was retrospectively found to be most impactful. Based on these findings, we propose that precise mapping of individual-specific brain network features could inform future interventions in patients.

**SIGNIFICANCE STATEMENT:** The network organization of every person’s brain is different, but non-invasive brain stimulation (NIBS) interventions do not take this variation into account. Here we demonstrate that the spatial positions of brain areas theoretically serving important roles in cognition, called hubs, differs across individual humans, but are stable within an individual upon repeated neuroimaging. We found that administering NIBS to these individual-specific hub brain areas impacted cognition more than stimulation of non-hub areas. This finding indicates that future NIBS interventions can target individual-specific, but cognitively-relevant features of human brains.

## INTRODUCTION

There is widespread interest in using non-invasive brain stimulation (NIBS) as a treatment for psychiatric and neurologic conditions (1), including addiction (2), obsessive compulsive disorder (3), stroke (4), and depression (5). The outcomes of these interventions, however, are variable across treated individuals. In the most widely adopted, Federal Drug Administration approved application of NIBS – repetitive transcranial magnetic stimulation (TMS) for the treatment of medication refractory major depression – only 29% percent of patients respond positively (6). Variation in patient response has been attributed in part to uncertainty regarding free parameters inherent to NIBS (7), including the stimulation site. Because the effects of NIBS are believed to propagate from the stimulation site in a manner constrained by the connectivity of the targeted brain area (8, 9), appropriate stimulation site selection is likely critical for therapeutic success, as it will determine whether or not stimulation effects spread throughout clinically-relevant neural circuitry.

Stimulation site selection strategies often focus upon anatomical landmarks (10, 11) and local tissue properties (12). The same anatomical area, however, exhibits different patterns of functional connectivity across individuals (13–16). In other words, the same stimulation site in different individuals is not necessarily functionally equivalent – which theoretically could contribute to the variable outcomes of NIBS interventions. The advent of techniques for precisely characterizing the areal (17, 18) and network organization (14, 15) of individual human brains has set the stage for the development of personalized NIBS protocols, which in principle can increase the likelihood of producing more consistent outcomes in patients (19).

A connectomics framework, in which brain areas (“nodes”) engage in networked communication within and across brain networks (“modules”), is theoretically well-suited for mapping an effective stimulation site on an individual basis. This approach can capitalize on the idea that a node’s role in a network can be inferred from its connections (20). Of particular interest are select nodes, termed “connector hubs” (hereafter referred to as hubs), connected to multiple modules and critical for the function of many networks found in nature (21). Evidence from biophysical models (22, 23) and lesions in stroke patients (24, 25) indicates that hub brain areas could serve outsized roles in the human brain as well. For this reason, we predicted that administering NIBS to hubs mapped in single-subjects using resting-state fMRI (rsfMRI) would impact cognition more than non-hubs. rsfMRI is increasingly considered for the purposes of non-invasively mapping stimulation sites in NIBS therapies (26–28), in part because it circumvents task-related confounds (29). If this prediction is borne out, it would implicate hubs as compelling candidate targets in NIBS interventions. In addition, it would provide the first causal evidence for the importance of hub brain areas mapped prospectively in individual human brains.

We first assessed the feasibility of mapping hubs using rsfMRI in single-subjects, and employing them as NIBS targets using the Midnight Scan Club (MSC) (16), a publicly-available dataset of highly-sampled individuals. We discovered that the spatial positioning of hubs is variable across individuals, but highly-reproducible within individuals when mapped using large quantities of rsfMRI data. In a prospective, double-blind, within-subject NIBS experiment we then mapped a hub brain area in twenty-four healthy participants. We predicted that administering an inhibitory form of TMS to hubs would disrupt information processing during cognition more than inhibition of a non-hub area on the same gyrus. To test this prediction, we fit a drift diffusion model to each participant’s performance on an N-back working-memory task after a hub and non-hub area was inhibited using TMS (counter-balanced administrations >24 hrs. apart). Although hub inhibition theoretically should impact multiple forms of cognition, working-memory was selected because it is associated with increased communication between segregated brain networks (30, 31) that is potentially facilitated by hub regions (32). Stimulation sites were constrained to right middle frontal gyrus to ensure both targets were in a brain region thought to be relevant for working-memory a priori (33).

## RESULTS

### Single-subject hub estimates are highly-reproducible

We estimated the degree to which discrete cortical areas (“parcels”) in each MSC participant function as hubs using the graph theory metric, participation coefficient (34). Nodes with higher participation coefficient values (“hubs”) have edges that are distributed across more network modules than those with lower values (“non-hubs”) (**Figure1A**). We sought to first determine the amount of rsfMRI data necessary for achieving reproducible single-subject hub estimates. Non-overlapping epochs were randomly sampled from each MSC participant’s 5-hour rsfMRI time-series. A set of participation coefficients for parcels with a centroid in right middle frontal gyrus was calculated separately using each epoch, and reproducibility quantified as the rank spatial correlation of the two sets. This procedure was repeated 10^3^ times per epoch, with epoch lengths ranging from 1 to 60 minutes, in 1-minute steps. Reproducibility was low when using commonly-utilized quantities of rsfMRI data (i.e., 5-10 minutes). With larger quantities of rsfMRI data, however, single-subject hub estimates were highly-reproducible (see **Figure1B** for MSC01 and **SI 1** for all participants). For example, we observed an average *r*_s_ of 0.82 ± 0.10 when using 45 minutes of rsfMRI data.

**Figure 1.**
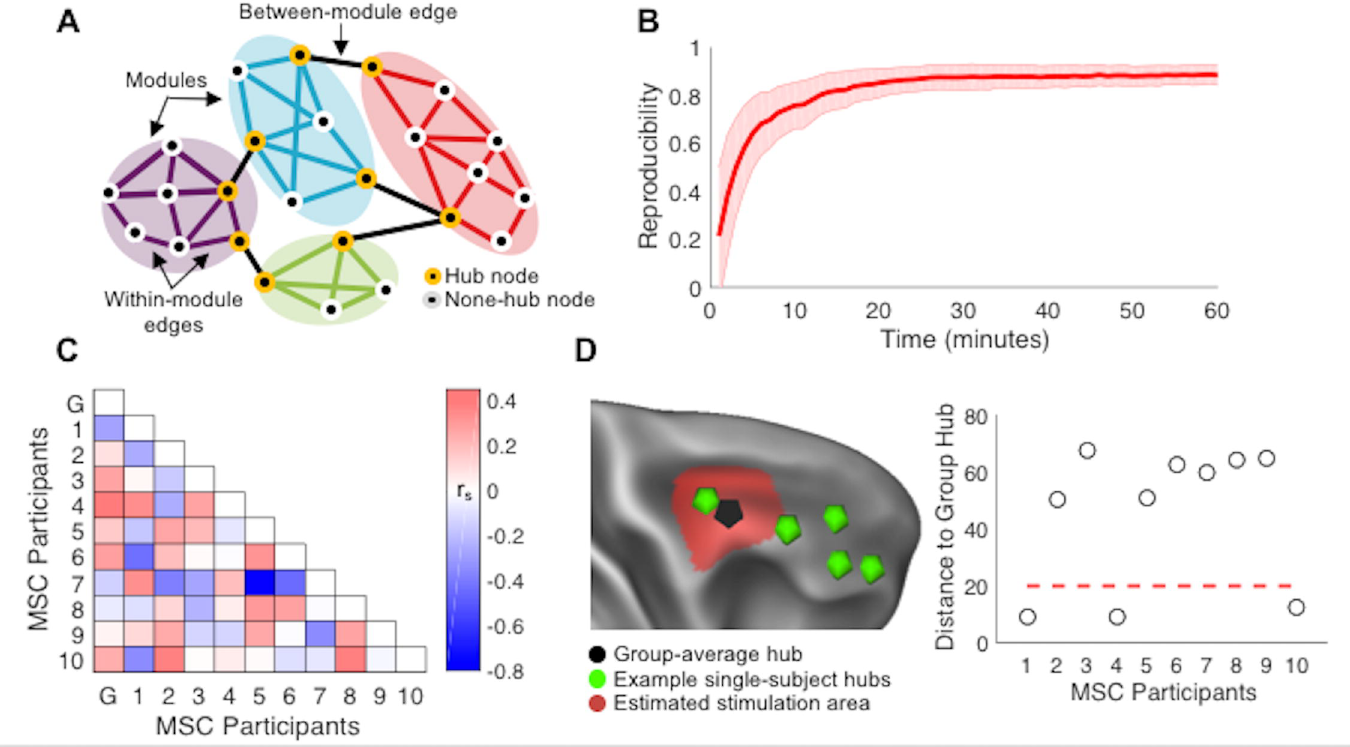
Hub nodes link multiple network modules, and have a high participation coefficient, whereas non-hub hub nodes have edges constrained within their network module, and have a low participation coefficient (**A**). Reproducibility of single-subject hub estimates improves with greater quantities of per-individual rsfMRI data in. Shaded red error denotes standard deviation (**B**). Similarity matrix summarizes inter-individual variation in spatial distribution of participation coefficients (**C**). Distances between example individual-specific hubs (green foci) and the group-average (“G”) hub. Red line and area of cortex surrounding the group-average hub (black foci) represents an estimated spatial resolution of TMS (**D**).

### Hubs are idiosyncratic features of functional brain organization

We next quantified inter-individual variation in the spatial distribution of hub estimates using pairwise rank spatial correlations (**Figure1C**). The spatial distribution of hub estimates across individuals was not similar (average *r*_s_ = −0.01 ± 0.26). Furthermore, single-subject hub estimates were not similar to their collective group-average (*r*_s_ = 0.09 ± 0.21). The practical implications of this finding for the proposed NIBS experiment was assessed by estimating the effective stimulation zone (**Figure1D**, red) surrounding the group-average hub (the highest participation coefficient parcel). Administering cTBS to this target would have theoretically failed to inhibit a hub in 70% of MSC participants (**Figure1D**), assuming a liberal spatial resolution of 0.5-2cm (35, 36). Thus, administering cTBS to a group-average hub might fail to impact hubs in some individuals. Collectively, these findings motivated the decision to map hub and non-hub stimulation sites on an individual basis using large quantities of rsfMRI data in the subsequent NIBS experiment.

### Precision mapping and inhibitory stimulation of hubs

Twenty-four healthy participants completed all three sessions (**Figure2A**). An automated pipeline (**Figure2B**) mapped a hub (the highest-participation coefficient parcel) and non-hub (the lowest-participation coefficient parcel ≥ 20mm from the hub) in the right middle frontal gyrus of each participant (**Figure2C**) using 45 minutes of rsfMRI. Native-image space coordinates corresponding to the hub and non-hub were then pseudo-randomly assigned as targets for two follow-up sessions (average interval between sessions = 5.8 ± 5.3 days). Immediately prior to performing an N-back fMRI task during these sessions, continuous theta burst stimulation (cTBS) (37) was administered offline and guided by neuronavigation. The aftereffects of cTBS are thought to last up to fifty-minutes (38), long-enough to complete the twelve-minute N-back task. Notably, we retroactively determined that hub and non-hub stimulation sites did not significantly differ in their anatomical positioning, nodal degree, baseline N-back activation, or distance to the stimulating coil (**SI 2**).

**Figure 2.**
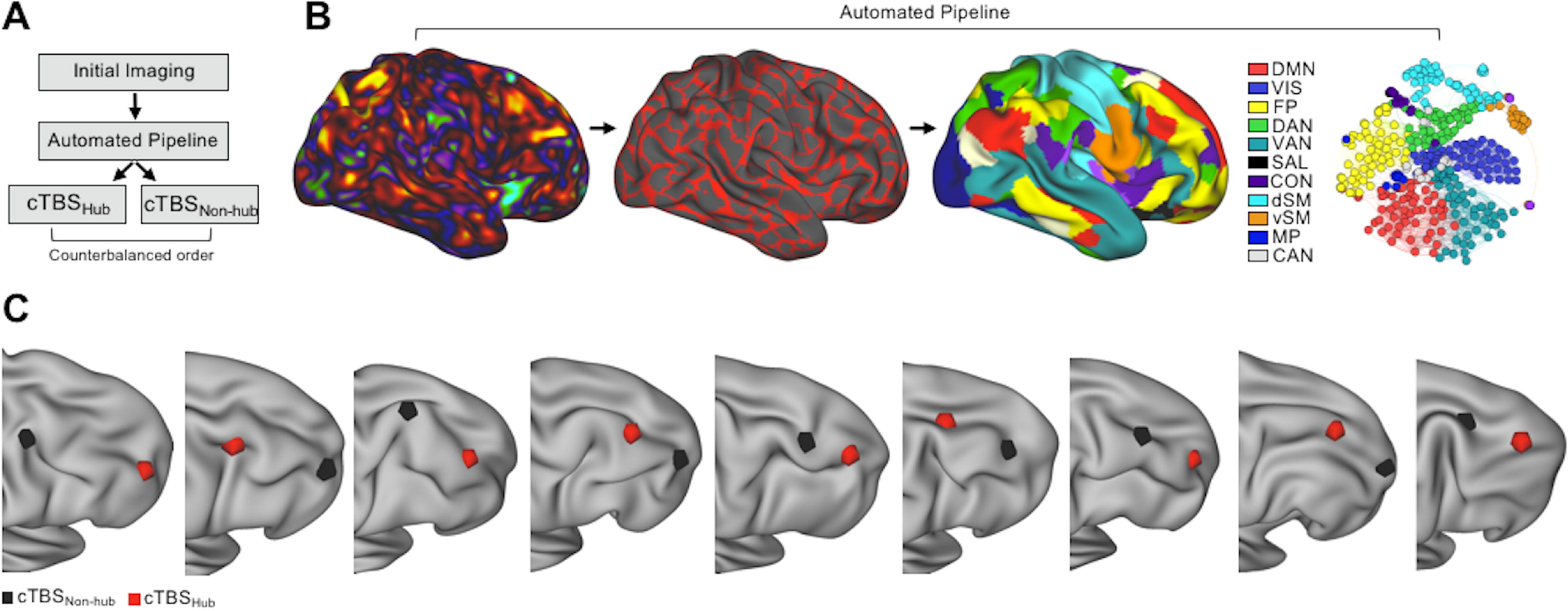
A prospective, within-subjects, double-blind experimental design (**A**). 45-minutes of rsfMRI and a high-resolution structural image collected during an initial study visit was submitted to an automated pipeline (**B**). Major pipeline steps include the construction of an individual-specific areal parcellation (red denotes boundaries between functional areas) and network structure (colors denote unique networks), and sparse functional connectivity matrices (a spring graph with a density of 1% for visualization). Participation coefficients for parcels with a centroid in right middle frontal gyrus were calculated. Parcels with the highest (“hub”) and lowest (“non-hub”) participation coefficients were pseudo-randomly assigned as targets for follow-up sessions [**C**: hubs (red) and non-hub (gray) cTBS targets in nine example participants]. DMN = default mode network, VIS = visual, FP = fronto-parietal, DAN = dorsal attention network, SAL = salience, VAN = ventral attention, AUD = auditory, dSM = dorsal somatosensory, vSM = ventral somatosensory, mPar = medial parietal, CAN = contextual association network, CON = cingulo-opercular.

### Hub inhibition disrupts information processing during working-memory

We tested our hypothesis that inhibiting hubs with cTBS would disrupt information processing using a drift diffusion model. This model takes the mean and variance of response times for correct trials, and mean accuracy as inputs and calculates cognitively-relevant latent variables indexing the rate of information processing (“drift rate”), response conservativeness (“boundary separation”), and non-decision time. Of these three diffusion parameters, drift rate was most relevant for our prediction that hub inhibition will disrupt information processing, as slower drift rates index worse N-back performance (39). Slower drift rates can be interpreted as slower, more variable response times and fewer correct responses (**Figure3A**).

**Figure 3.**
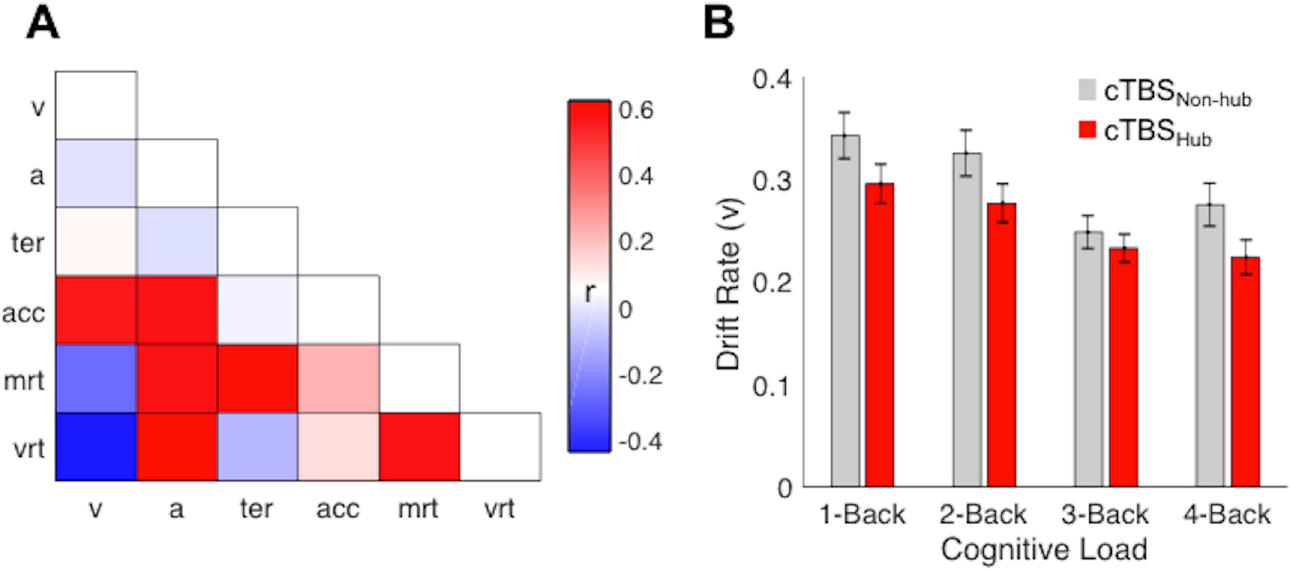
Drift diffusion modeling of N-back performance following hub and non-hub inhibition with cTBS. A correlation matrix, where entries denote the pairwise relationships of input (mrt = mean response time, vrt = variation in response time) and output (v = drift rate, a = response boundary, ter = non-decision time, acc = mean accuracy) parameters, was constructed to aid in the interpretation of diffusion parameters (**A**). A 2 ^x^ 4 repeated measures ANOVA (target ^x^ load) performed on drift rates revealed that hub cTBS disrupted drift rates more than non-hub cTBS (**B**).

A 2 ^x^ 4 repeated measures ANOVA (target ^x^ load) performed using drift rates revealed a main effect of target [*F*(1,23) = 8.60, *p*_bonferroni_ = 0.02], such that drift rates following hub inhibition were slower than those after non-hub inhibition (**Figure3B**). This finding demonstrates, as predicted, that hub inhibition disrupts information processing more than inhibition of a nearby non-hub. Notably, this effect remained in ANCOVA models where differences in anatomical positioning, baseline activation, and distance to TMS coil were included as a between participant covariate (**SI 3**). As expected, we also observed a main effect of load [*F*(1,23) = 31.50, *p*_bonferroni_ < 0.001], confirming that higher cognitive loads were more difficult than lower loads. Target and load did not interact [*F*(1,23) = 0.10, *p*_bonferroni_ = 1.00], however, indicating that the difference in drift rate following hub and non-hub inhibition did not vary significantly by cognitive load. ANOVA models were also performed on the response boundary and non-decision time parameters (**SI 4**).

### Hubs may belong to discrete, functional-relevant categories

Our experimental design assumed that hubs belong to a single category of objects with global connections, akin to a “hub-spoke” network. Recent evidence, however, indicates that hubs instead tend to link discrete, non-random subsets of brain networks, and can be clustered using their cross-network profiles of functional connectivity into three categories (40). The three hub categories are termed according to their putative information processing roles – external, internal, and control (**SI 5**). We hypothesized post-hoc that the degree to which hubs targeted with cTBS in the present investigation resembled these hub types could account for variation in drift rate change (drift rate_hub_ – drift rate_non-hub_) averaged across cognitive loads. We tested this possibility in an exploratory analysis, using a stepwise linear regression model with hub type resemblances (Fisher-transformed correlation coefficients) as predictors. Resemblance with the external hub type was the only significant predictor (β = −0.41, ΔR^2^ = 0.17, *p* = 0.04). The external hub type is distinguished by linking control networks, including the cingulo-opercular network and dorsal attention network, with processing systems thought to represent external information, including the visual and somatomotor networks.

## DISCUSSION

We present three findings in this investigation. First, cortical hubs mapped using large quantities of per-individual rsfMRI are highly-reproducible, idiosyncratic features of functional brain organization. Second, inhibiting a hub area with cTBS impacted working-memory performance, as measured using a drift diffusion model, more than inhibiting a non-hub area of the same gyrus. Third, inhibiting hubs linking select control-related networks with those relevant for processing external stimuli was most impactful. The theoretical and potential translational implications of these points are considered below.

### On intra-and inter-individual variation in single-subject hub estimates

The reproducibility curves for single-subject hub estimates reported here began low and rapidly reached asymptote, resembling sigmoid functions reported in similar analyses (41). Measurement error, due to the amount of per-individual data utilized, is thought to be a primary source of intra-individual variance in rsfMRI functional connectivity estimates. Other studies have also reported that small quantities of per-individual rsfMRI data fail to reliably characterize functional brain organization (16, 41, 42), likely due to the poor signal-to-noise ratio in BOLD fMRI data (43). Consistent with this perspective, we found that large quantities of rsfMRI data are needed for reproducible single-subject hub estimates. It is noteworthy that Gordon and colleagues (16) reported that whole-brain hub estimates were less reproducible. Our analysis, however, was constrained to right middle frontal gyrus. Thus, this difference could be explained by less intra-individual variation in rsfMRI functional connectivity estimates in this brain area (13, 44).

The practical significance of inter-individual variation in topological features functional brain organization, including hubs, for future work will depend on whether precision is necessary for the context at hand. Consider, for example, that the average distance between single-subject and the group-average hubs was on the order of centimeters. Thus, a group-average map may reflect a central-tendency in the spatial positioning of hubs sufficient for purposes of cartography (45), but interventions employing high-resolution techniques, such as TMS (35, 36), could benefit from targeting individual-specific hubs. With this concern in mind, we mapped hubs on an individual basis in our NIBS experiment.

### Interpreting the drift diffusion model parameters

The computational model applied in our investigation assumes an accumulation of noisy information supporting a decision process - whether or not the current stimulus matches one occurring *N*-trials ago. Conceptually, the drift rate parameter indexes the average amount of information accumulated per unit of time during this decision process (46). It is noteworthy that changes in drift rate, despite being conceptualized in terms of speed (i.e., faster or slower), are due to changes in both accuracy and response time distributions, such that slower drift rates reflect slower, more variable, less correct responses. Slower drift rates after hub inhibition could result from a disruption in the integration of task-relevant information distributed amongst brain networks bridged by the inhibited hub (39). This scenario could explain why inhibiting some hubs disrupted information processing more than others (i.e., if task-relevant information was represented in a subset of discrete brain networks). While working-memory was identified a priori as one form of complex cognition theoretically involving hubs, we believe that it is unlikely that hubs subserve any single cognitive process. Instead, the extent to which hubs are functionally specialized is more likely with respect to broader domains of cognitive processing (e.g., externally-vs. internally-oriented cognition) (40).

### Inhibition of different hub types differentially affects behavior

The network neuroscience literature generally conceptualizes hubs as a single class of nodes with uniform function and global connections (47, 48). Consistent with this perspective, our NIBS experiment was designed to inhibit a hub in each individual, without explicitly considering which networks it bridges. By retrospectively comparing hubs to independently identified categories of hubs (40), however, we assessed whether the variation in the behavioral effects of hub inhibition could be related to differences in cross-network connectivity. The degree to which hubs resembled a so-called external hub type was a significant predictor. Although speculative, the topological positioning of the external hub type, at the intersection of select control and sensorimotor networks, is well-suited for enabling information processing during a visual working-memory task requiring a motor response. An item maintained in working-memory is theoretically represented broadly in sensorimotor cortex and influenced by signals from control-related brain areas (49, 50). Thus, it is tempting to conclude that hub types could be specialized for specific forms of information processing. In principle, a future NIBS experiment could test this possibility empirically by prospectively mapping different hub types within an individual and testing for a dissociation.

### Implications for translational network neuroscience

Network theory is a powerful and increasingly widespread conceptual framework in neuroscience (51, 52). This approach distills the complexity of the brain into simpler mathematical representations (53), which in turn allows investigators to form and test tractable hypotheses regarding how a brain might process information in a networked fashion. Validating the core predictions of this framework is a necessary step towards establishing its translational value (54, 55). One of the most useful predictions in network neuroscience is that the role a brain area can be inferred from its connections (20). From this perspective, hub nodes should be more important than more peripheral nodes for network function. Evidence supporting this prediction to date in the human brain has come from biophysical models (22, 23) or lesions in stroke patients (24, 25), with no precise, causal manipulations of hubs mapped prospectively in single-subjects performed until the present investigation. Thus, beyond providing evidence for the importance of hub brain areas, this investigation highlights how network neuroscience could be leveraged in the future to inform personalized interventions in humans (56), including NIBS (57), but also neurofeedback or rehabilitation.

### Implications for rsfMRI-guided NIBS interventions

There is growing interest in using rsfMRI to guide targeted NIBS therapies (26–28). This is because rsfMRI can efficiently characterize the intrinsic functional network organization of a brain that is present across many task states (58) but is not limited by task-related confounds (29). We found that inhibiting hub and non-hub brain areas, which differed only in their rsfMRI connectivity, produced significantly different behavioral outcomes, despite being separated by only a few centimeters on the same gyrus. It is noteworthy that NIBS investigations generally use either a sham condition or an active control that is anatomically distinct. Thus, utilizing a nearby non-hub as an active control is a highly rigorous element of our experimental design (59). The success of this approach highlights the precision at which rsfMRI and select NIBS techniques may be able to map and manipulate functionally discrete areas of cortex in patients. Finally, future work can evaluate whether NIBS interventions could benefit from stimulating hubs mapped using rsfMRI in single-subjects. This strategy would be a significant departure from existing proposals for network-centric targeting (28, 60), as hubs are positioned at the intersection of multiple networks, which could afford an opportunity to modulate a wider range of symptoms. This strategy is conceptually well-suited for populations, such as depression (61), with heterogeneous clinical profiles and abnormal functional connectivity between multiple brain networks (62). Recently engineered TMS mini-coils (63) may afford an opportunity to test this hypothesis in a rodent model of treatment-resistant depression (64).

## CONCLUSIONS

This investigation joins an emerging field of precision connectomics (16, 41, 65) treating idiosyncrasies in functional brain organization as neurobiologically informative - and not noise. Our findings highlight how precise mapping of topological features of interest in single-subject connectomes could be leveraged in the future to guide interventions in the human brain. NIBS therapies may be particularly well-suited to adopt this approach.

## METHODS

### Midnight Scan Club

#### Participants

The Midnight Scan Club (MSC) dataset (16) was downloaded from OpenfMRI.org (ds000224). This dataset is comprised of ten participants aged 24-34 years (mean age = 29.1 years ± 3.3, 5F/5M) that underwent a total of 5-hours of rsfMRI (10×30 minutes, each session acquired at midnight on subsequent days). Further details regarding data acquisition and sample demographics are reported by (16). Details regarding analyses performed on the MSC dataset are included in the **Supplementary Information**.

### Non-invasive brain stimulation experiment

#### Participants

Twenty-four participants aged 18-28 years (mean age = 20.5 years ± 2.5, 11F/13M) were recruited from the Georgetown community after complying with the consenting guidelines of the Georgetown University IRB. Participants were screened for history of neurologic and psychiatric conditions, epilepsy, contraindications for MRI, and use of medications that increase likelihood of side effects following TMS.

### Data acquisition

All data was acquired on a Siemens Trio 3T with the participant’s head immobilized using head cushions. A high resolution structural T1 scan was acquired with the following parameters: MPRAGE: TR/TE = 1900/2.52 ms, 90-degree flip angle, 176 sagittal slices with a 1.0 mm thickness. Functional echo-planar images were acquired with the following parameters during each imaging sessions: 3 mm isotropic resolution, TR = 2000 ms, TE = 30 ms, flip angle = 90°, FOV = 192 × 192 mm. A T1 and three resting-state runs, each lasting 15-minutes and acquired successively, were collected during the initial imaging visit. An N-back task consisting of 20 blocks (5 blocks each of 1-, 2-, 3-, and 4-back loads, in pseudo-randomized order) was administered during each study visit. Blocks consisted of 9 letters, each presented for a duration of 500ms and with an inter-trial interval of 1500ms. Participants were instructed to provide a right-hand button press for targets and a left-hand button press for non-targets as quickly and accurately as possible. Of the 180 trials, 32 were targets, and either one or two targets were in any given block. Stimuli were presented on a back-projection screen using the E-Prime software.

### Data preprocessing

Functional images were corrected for differences in motion and slice timing acquisition, and co-registered into each participant’s anatomical image using SPM12 (Wellcome Department of Cognitive Neurology, London, United Kingdom). Functional data was denoised using the aCompCor strategy in the CONN toolbox. Denoising steps included linear de-trending and nuisance regression (5 principle components from white matter and cerebrospinal fluid masks from an MPRAGE segmentation; 6 motion parameters and first-order temporal derivatives; and point-regressors to censor time points with mean frame-wise displacement > 0.2mm). Residual time-series were band-pass filtered (0.01 Hz < *f* < 0.1 Hz). Temporal masks were created to flag motion-contaminated frames for scrubbing. Contaminated volumes were identified by frame-by-frame displacement (FD) calculated as the sum of absolute values of the differentials of the 3 translational motion parameters and 3 rotational motion parameters. On average, 76 ± 3% of the rsfMRI time-series was retained after motion censoring.

### Surface file generation

Following volumetric co-registration, white and pial anatomical surfaces were generated from each participant’s native-space MPRAGE using Freesurfer’s recon-all pipeline (version 5.0). The fsaverage-registered left and right hemisphere surfaces were then brought into register with each other in fs_LR space (66) and resampled to a resolution of 32k vertices using Caret tools. Denoised fMRI time-series within the cortical ribbon were mapped onto each individual’s midthickness surface and spatially smoothed (σ=2.55). Both left and right surfaces were combined into the Connectivity Informatics Technology Initiative (CIFTI) format using Connectome Workbench (67), yielding time courses representative of the entire cortical surface, excluding non-gray matter tissue, and sub-cortical structures.

### Automated pipeline for identifying stimulation targets

All of the steps below were performed in a single automated pipeline, allowing both the cTBS administrator (CJL) and participant to remain blinded. First, a boundary-based areal-parcellation (18) was generated using each participant’s denoised resting-state CIFTI dataset. Parcels were generated using the watershed by flooding procedure. Parcels smaller than 10 vertices (~20mm^2^) were removed. The number of resultant parcels varied between individuals (mean parcel count: 548, range: 473-603), consistent with the number of parcels observed in other single-subject areal-parcellations (16, 41). Second, parcels were assigned to a one of twelve canonical networks, defined in an independent sample of healthy adults (N=120) using the InfoMap algorithm (68), via a template matching procedure (14). Finally, the temporal correlation between each parcel and all other parcels was computed using the denoised motion-censored rsfMRI time-series, yielding a parcel ^X^ parcel functional connectivity matrix for each participant. Local connections (those < 30mm in geodesic space) were eliminated, to avoid local blurring of signals between adjacent parcels. The participation coefficient and nodal degree was calculated for each parcel. To do so, we first calculated ten participation coefficients and degree values for each parcel using binarized functional connectivity matrices, each with a unique density (1-10%, in 1% steps), following other investigations (69). The final participation coefficient and degree for each parcel was the average of these ten values. Participation coefficient was calculated using the equation below, where *M* is the total set of networks, k_*i*_ is the number of edges associated with node *i*, and k_*i*_ (*m*) is the number of edges between node *i* and all nodes in network *m*.

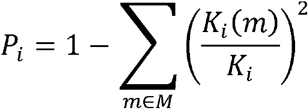

Because the participation coefficient of a node with few edges is dubious, we eliminated from consideration parcels with a degree in the bottom quartile of whole-brain values. Remaining parcels with a centroid falling within right middle frontal gyrus, defined using Freesurfer gyral labels, were identified. The hub was defined as the parcel with the highest participation coefficient value. The euclidean distance between the hub and all other parcels in right middle frontal gyrus was calculated. The non-hub was defined as the parcel with the lowest participation coefficient value >20mm from the centroid of the hub parcel in euclidean space. This minimum distance was enforced to enable selective targeting of the two parcels, given evidence that the spatial resolution of TMS is 0.5-2cm (35, 36). Native-space MPRAGE coordinates for the hub and non-hub were pseudo-randomly assigned to the two follow-up sessions.

### Continuous theta-burst stimulation and MRI-guided neuronavigation

cTBS was applied at 80% of active motor threshold using a MagPro x100 device (MagVenture, Inc., Atlanta, GA) with a passively coiled MCF-B70 figure 8 coil. cTBS is a safe (70) and inhibitory form of patterned TMS involving three 50Hz pulses in trains repeated at 200-ms intervals (37). Parcel centroids were targeted using the Brainsight 2 Frameless stereotactic system for image guided TMS research (Rogue Research, Montreal, Canada). This system uses infrared reflectors attached to a headband worn by the subject to co-register the MPRAGE with the participant’s head. The coil was co-registered via infrared reflectors. Active motor threshold was defined as the intensity required to induce a motor evoked potential in the contralateral FDI muscle when pulses were applied to right motor cortex during a mild sustained contraction. Muscle twitches were measured using surface electrodes placed on the FDI muscle, which are connected to an electromyography device incorporated into the BrainSight system.

### Drift diffusion modeling of task performance

A drift diffusion model is a well-validated computational model that is advantageous to considering response times and accuracy in isolation from one another (71). We selected an EZ-diffusion model (71), in place of alternative diffusion fitting routines, as it is more effective in resolving individual differences in parameter values (72) and does not require characterizing the distribution of incorrect response times, making it well-suited for use in populations where the error rate is relatively low. An EZ-diffusion model does not perform well if there are contaminants (e.g., an extremely fast response, indicative of a guess) present (73). For this reason, we removed trials with a response time <300ms from consideration. A 2 ^x^ 4 (target ^x^ load) repeated measure ANOVA was performed on each of three diffusion model parameters - drift rate, boundary separation, and non-decision time. Tables containing summary statistics for diffusion model inputs can be found in **S6** & **S7**. Reported p-values were bonferroni corrected, to account for the three ANOVA models that were performed.

### Clustering hubs into discrete subtypes

We calculated the average functional connectivity between hub parcels and all parcels of each brain network that were >30mm in geodesic space. The similarity of these resultant hub cross-network connectivity profiles to three hub type templates was calculated (details regarding the creation of the hub type templates is described in **S5**). This resulted in a 24 ^x^ 3 (participant ^x^ hub type) array of Fisher-transformed correlation coefficients. A stepwise linear regression analysis was performed using these coefficients as predictor variables and the change in drift rate (drift rate_hub_ – drift rate_non-hub_) averaged across loads as a dependent variable. Note that we used stepwise linear regression because the average hub type profiles were not independent from one another. Stepwise linear regression was performed using the Matlab function (“stepwiseglm”).

## Acknowledgements

This work was supported by Dean Toulmin’s Pilot Project Award from Georgetown University Medical Center to PET and CJV. We would like to thank the research staff at the Center for Functional and Molecular Imaging and Junaid Merchant for their assistance with data collection.

